# Loopsim: Enrichment Analysis of Chromosome Conformation Capture with Fast Empirical Distribution Simulation

**DOI:** 10.1101/2024.06.03.595407

**Authors:** Gideon Shaked, Haihan Zhang, Zhaolin Zhang, Johann E. Gudjonsson, James T. Elder, Matthew T. Patrick, Lam C. Tsoi

## Abstract

**Summary:** Gene regulation is intricately influenced by the three-dimensional organization of the genome. In particular, chromatin can exist in loop structures that enable long-range regulatory interactions. By utilizing chromosome conformation capture techniques such as Hi-C, valuable information regarding the organization of these loop structures in 3D space can be obtained. While functional/feature enrichment has become a standard downstream analysis for different genomic data to provide biological context, tools that developed specifically for high throughput assays capturing chromosome conformation are relatively limited. Here, we present Loopsim, a command-line application that performs enrichment analysis on Hi-C loop profiles against user-defined regions. Loopsim efficiently simulates a background distribution using a distinctive sampling approach that considers loop size, intervals, loop-loop distances, and structure; it then computes loop-level statistics based on the empirical null distribution.

**Availability:** Loopsim is a Python package available via PyPI (https://pypi.org/project/loopsim) and the source code is available on GitHub (https://github.com/CutaneousBioinf/Loopsim) under the MIT license.

## 1 Introduction

The organization of the genome plays a crucial role in regulating gene transcription, with chromatin loops connecting distant genomic sites to facilitate spatial proximity in three-dimensional space (Kadauke and Blobel, 2009). Chromosome conformation capture methods, such as Hi-C, can generate high-quality data for chromatin organization (Berkum, et al., 2010). Multiple high throughput chromosome conformation methods including promoter capture Hi-C, ChIA-PET, HiChIP, and PLAC-seq have been developed to identify loops associated with gene regulation (Durand, et al., 2016; Roayaei Ardakany, et al., 2020; Wolff, et al., 2022). These chromosome conformation capture methods have significant implications in uncovering the mechanisms of gene regulation, including the interpretation of genetic signals, and revealing the structure of the regulatory complex as well as the gene targets of enhancer regions (Sahlén, et al., 2021; Shi, et al., 2021). Current downstream analyses include chromatin loop and topologically associated domain identifications, as well as differential loop analysis and chromatin compartment separations (Han and Wei, 2017). However, despite the present success in functional enrichment approaches developed for different genomics and epigenomics platforms (Huang, et al., 2009), there are limited tools that integrate functional annotations to provide the biological meanings of loops identified in Hi-C experiments.

In this study, we present Loopsim, a tool designed to detect the statistical significance of overlaps between identified chromatin loops and genomic regions of interest (e.g. GWAS signal; regulatory regions identified in ChIP-seq or ATAC-seq). In contrast to existing tools (**Table 1**), Loopsim is packaged as a pipeline for enrichment analysis that includes Hi-C data validation, simulation, and visualization. It simulates the empirical null distribution of Hi-C loop profiles as background for enrichment analysis. The background simulation algorithm takes into consideration factors including loop size, intervals, loop-loop distances, and structural characteristics, while also implementing logic to avoid succumbing to bias introduced as a result of particular characteristics of the inputted Hi-C loop profile.

**Table 1.**
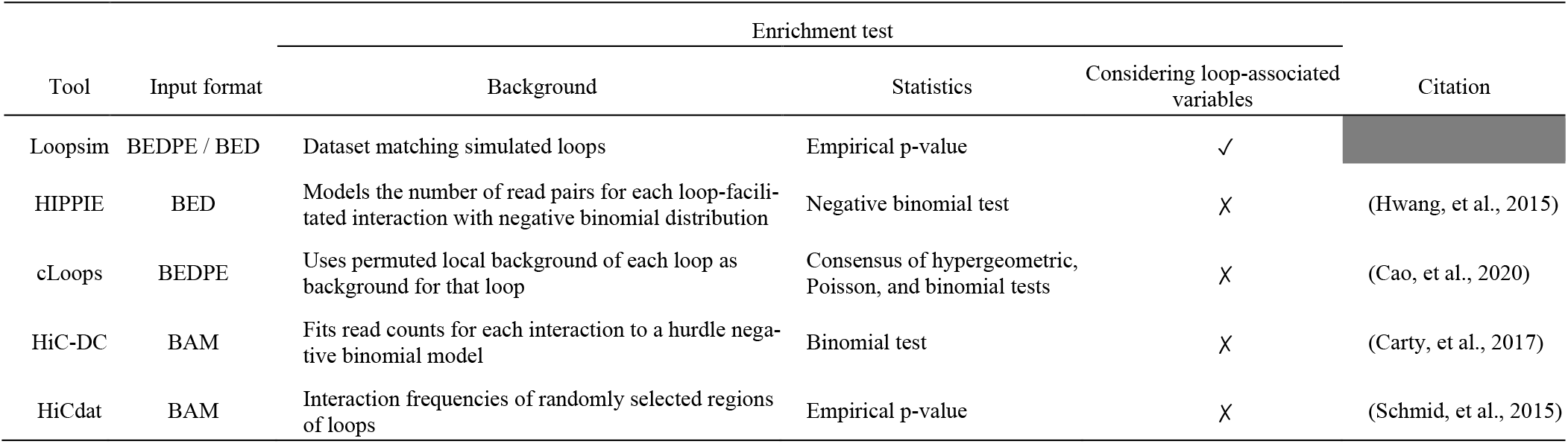
Comparison of Loopsim with existing chromosome conformation capture enrichment analysis tools.

## 2 Implementation and Features

Loopsim is a command line program capable of working with Hi-C loop profiles, which are loop-calling results from standard Hi-C data produced with tools such as HICCUPS (Rao, et al., 2014) or Mustache (Roayaei Ardakany, et al., 2020). Loopsim provides a pipeline (**Figure 1**) that comprises input data preprocessing, identification of chromatin loops whose end regions overlap with genomic regions of interest, simulation of a background distribution of Hi-C loop profiles, and enrichment analysis of the original Hi-C loop profile against said background distribution. Furthermore, Loopsim is flexible concerning the data it can process, in that it can handle an arbitrary set of chromosomes and associated loci information.

**Fig 1.**
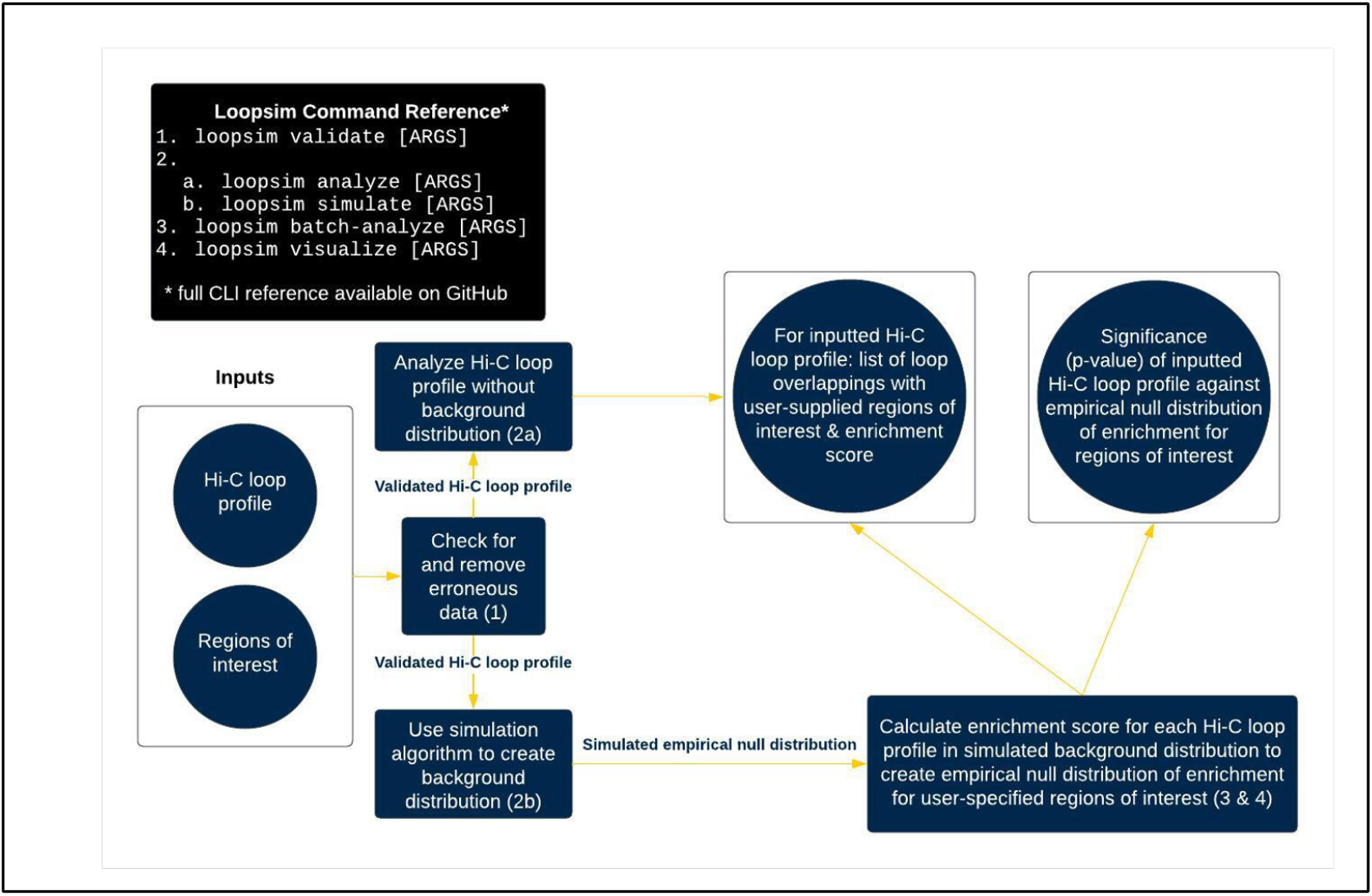
Loopsim pipeline

### Validation

Prior to running the rest of the pipeline, Loopsim performs quality control on the input Hi-C loop profile, including validation checks to exclude chromatin loops that start and end on different chromosomes (Kadauke and Blobel, 2009) and filter out unrealistically long loops (e.g. ≥100 kb by default (Jackson, et al., 1990)); these values are user-configurable. Loopsim also checks and performs filtering for chromatin loops with differently sized end regions and for overlapping end regions, both conditions potentially caused by data processing errors. After all checks are complete, Loopsim sorts the chromatin loops of the Hi-C loop profile by chromosome and the start locus of the loop.

Next, Loopsim can be used to generate a simulated empirical background distribution of Hi-C loop profiles against which to conduct enrichment analysis (**Figure 1**); alternatively, Loopsim can be used to perform a cursory analysis of the inputted Hi-C loop profile.

### Cursory analysis of a Hi-C loop profile without simulated empirical background distribution

Loopsim calculates an “enrichment score” for the input Hi-C loop profile, denoting the proportion of chromatin loops that overlap with genomic features of interest while also providing a list of the overlapping loops. Such information would inform a user, as a preliminary analysis step, whether their Hi-C data contains loops with potential for significant genomic interactions.

### Enrichment analysis of a Hi-C loop profile with a simulated empirical background distribution

Using the method described in **Section 3**, Loopsim generates a simulated empirical background distribution of Hi-C loop profiles to a user-specified size. Then, Loopsim performs a one-tailed difference of means test, comparing the average proportion of chromatin loops overlapping regions of interest in the empirical null distribution with the proportion of overlapping loops in the input Hi-C loop profile. Loopsim employs this test to ascertain if the real Hi-C loop profile exhibits a significantly higher proportion of chromatin loops with ends overlapping genomic regions of interest compared to the background distribution, and the result of this test is a measure of the enrichment of the original Hi-C loop profile. Loopsim can also compute the enrichment statistic parametrically, and a normality test is provided.

## 3 Simulation Algorithm

In order to generate a simulated background distribution of Hi-C loop profiles, Loopsim employs the following partially stochastic procedure (**Figure 2**) to simulate an individual Hi-C loop profile, and simply repeats it a user-specified number of times. To create one simulated Hi-C loop profile, Loopsim iterates loop by loop through the original Hi-C loop profile and generates corresponding loops for the simulated Hi-C loop profile. To generate each simulated chromatin loop, Loopsim uses one of two chromatin loop simulation methods, *random simulation* or *nonrandom simulation*. For Loopsim to perform *nonrandom simulation*, the distance between the real chromatin loop and the previous real chromatin loop must be less than 1 Mb. This condition exists under the assumption that loci within 1 Mb of each other are more likely to interact with each other (Pennacchio, et al., 2013). As such, we want to retain such arrangements from the real chromatin loops when generating simulated chromatin loops. In *nonrandom simulation*, Loopsim first measures the distance between the first ends of the current and preceding real chromatin loops. It then copies the previous simulated chromatin loop and positions this copy the same number of base pairs ahead of the previous simulated chromatin loop. In the opposite case that the distance between the real chromatin loop and the previous real chromatin loop is greater than or equal to 1 Mb, Loopsim performs *random simulation*, where it copies the previous real loop and positions the copy at a random locus within the same chromosome.

**Fig 2.**
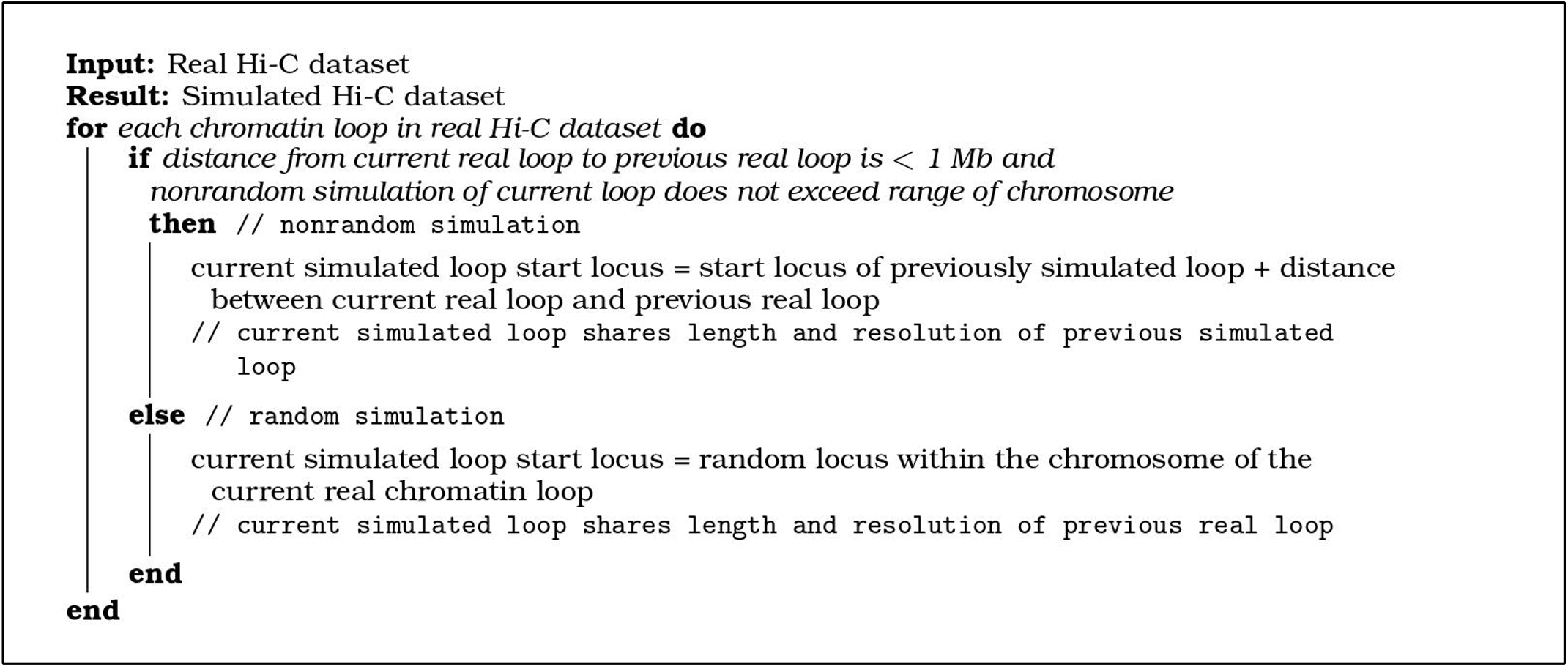
Pseudocode for Loopsim simulation algorithm

## 4 Results

**Figure 3A** presents an example output from our Loopsim program. To evaluate Loopsim’s performance, we used a Hi-C loop profile generated by deep sequencing Hi-C libraries (~1 billion reads/reaction). Then, using a contact map combining resolutions of 5Kb and 10Kb (resolution in this case meaning the length of the interval of either loop end), we identified Hi-C loops in each library using Mustache (Roayaei Ardakany, et al., 2020). The Hi-C loop profile was enriched for GWAS loci for psoriasis (Dand, et al., 2023). We ran Loopsim on this Hi-C loop profile and created simulated Hi-C background distributions of various sizes.

**Fig 3.**
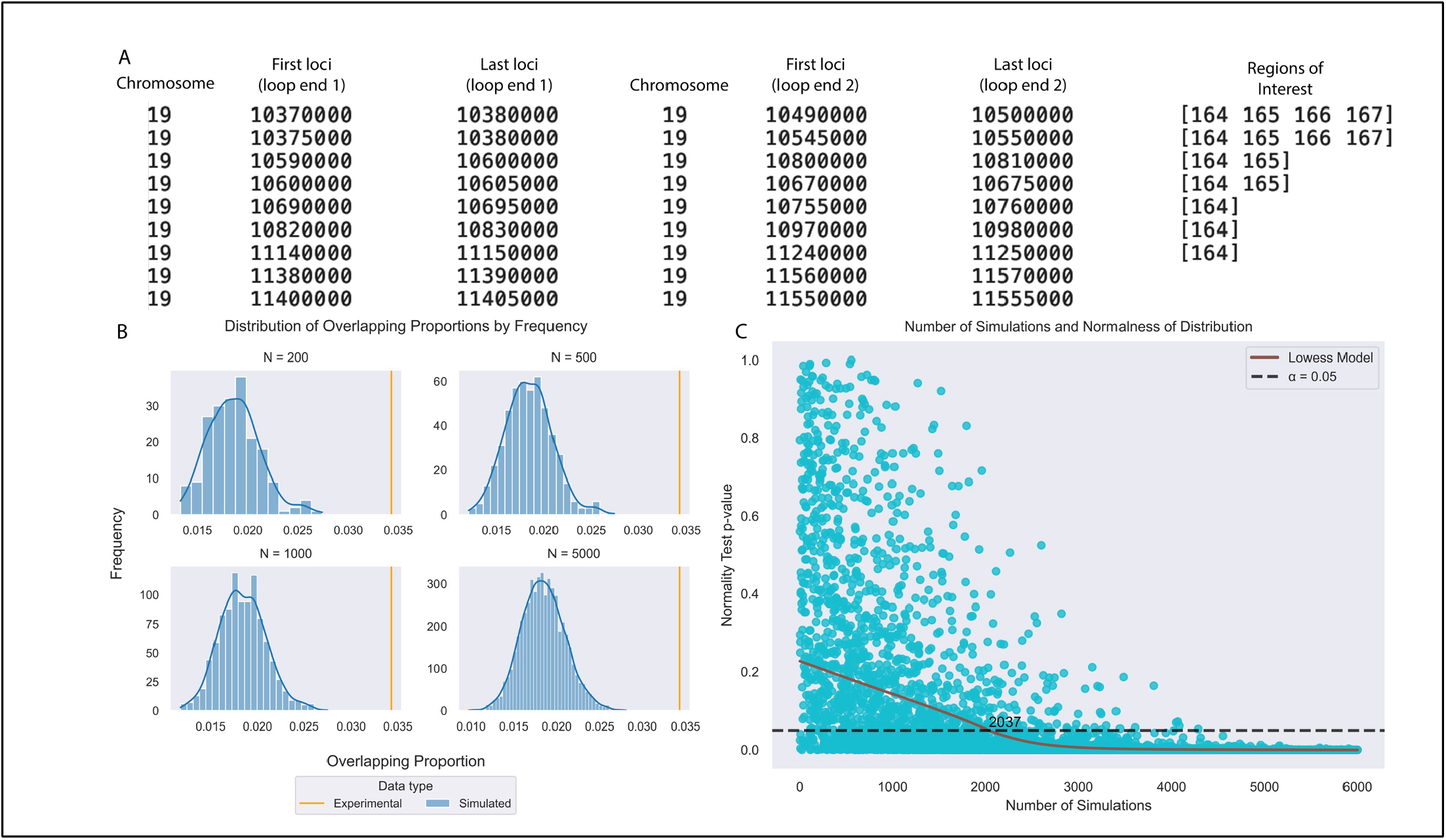
(A): Simulated Hi-C background distributions of different sizes were generated with Loopsim. For each simulation in each distribution, the proportion of enriched chromatin loops (those overlapping genomic regions of interest) was calculated. The proportion of enriched chromatin loops was also calculated for the corresponding experimentally produced Hi-C loop profile. (B): A D’Agostino-Pearson test of normality was performed on varying sizes of randomly sampled Hi-C background distributions. A locally weighted linear regression model was formed to indicate how the typical normality of Loopsim’s simulated Hi-C distributions varies with the number of simulated Hi-C loop profiles in the distribution. (C): Part of the output of the Loopsim analysis as demonstrated in **Section 4**. The first three columns from the left represent the first end of each chromatin loop, representing the chromosome, first loci, and last loci, respectively. The second three columns represent the last end of each chromatin loop in the same format. The last column represents the regions of interest that each chromatin loop overlaps with. For example, the loop on the second row overlaps with regions 164, 165, 166, and 167, whose details are represented in a separate file.

Relative to the simulated background distribution, the experimentally derived Hi-C loop profile showed a higher degree of enrichment for psoriasis loci, as shown by the experimental data having a greater overlapping proportion (**Figure 3B**). This observation underscores the accuracy of Loopsim in simulating an appropriate background distribution. Furthermore, we observed that Loopsim has a runtime approximately lin-early proportional to the number of Hi-C loops in the dataset. We can also see that after approximately 2,000 simulations, Loopsim’s simulated background distribution became approximately normal. We suggest that the user must specify at least approximately 2,000 simulations to generate a statistically normal, realistic simulated background distribution (**Figure 3C**).

## 5 Conclusions

We developed Loopsim to allow researchers to generate an accurate simulated empirical background distribution for Hi-C data analysis by considering loop size, intervals, distances, and structural characteristics. Applied to Hi-C data enriched for psoriasis, Loopsim successfully identified significant chromatin loops overlapping with GWAS loci, showcasing its potential for revealing meaningful genomic interactions. Loopsim’s user-friendly interface, comprehensive data validation, and robust end-to-end analysis pipeline make Loopsim an asset to genomic research. By enabling better enrichment analysis of Hi-C data, Loopsim can be used to enhance our understanding of 3D genome organization and gene regulation.

